# Initiating mammalian reproduction from a single male genome

**DOI:** 10.64898/2026.08.03.742506

**Authors:** Shogo Matoba, Nonoko Umeda, Mizuki Sakamoto, Sota Komatsubara, Dinh Quoc Pham, Asako Okamoto, Aoi Ito, Takumi Ando, Hirosuke Shiura, Kimiko Inoue, Atsuo Ogura, Toshiaki Hino, Takashi Ishiuchi

**Affiliations:** Integrative Developmental Engineering Division, RIKEN BioResource Research Center, Tsukuba, Ibaraki 305-0074, Japan; Cooperative Division of Veterinary Sciences, Tokyo University of Agriculture and Technology, Fuchu, Tokyo 183-8509, Japan; Faculty of Life and Environmental Sciences, University of Yamanashi, Yamanashi 400-8510, Japan; School of Biotechnology, International University, Vietnam National University, Ho Chi Minh 70000, Viet Nam; Department of Biological Sciences, Asahikawa Medical University, Asahikawa, Hokkaido 078-8510, Japan; Agro-Biological Resource Sciences, University of Tsukuba, Ibaraki 305-8577, Japan

## Abstract

Sex in mammals is unidirectionally determined by the configuration of sex chromosomes. In contrast, several species, including fishes, can switch sexes in response to environmental cues, illustrating the reproductive advantages of sexual plasticity. Thus, technologies that can bypass the unidirectional nature of mammalian sex determination could open new avenues in reproductive biology. Here, by establishing a method that enables robust Y-chromosome elimination in embryos, we provide a strategy to regulate sex in mice. We show that this approach efficiently induces male-to-female sex reversal by releasing the Y chromosome as micronuclei during early development. When integrated with optimized somatic cell nuclear transfer, it enabled the parallel production of both male and sex-reversed female mice from a single male genome source. Offspring were successfully obtained from crosses between these male and sex-reversed female clones, demonstrating that this “dual-sex cloning” strategy can initiate sexual reproduction solely from a male somatic genome. These findings highlight dual-sex cloning as a valuable platform for securing mammalian reproduction and biodiversity from limited genetic resources.

## Main Text

Reproduction is the most fundamental task that organisms must accomplish. To achieve this, many organisms have adopted sexual reproduction, which requires both female and male individuals. In mammals, sex is determined unidirectionally by the presence or absence of the Y chromosome, with XX individuals developing as females and XY individuals as males. Consequently, a skewed sex ratio in a population can threaten reproduction, as seen in endangered animals. In contrast, some fish species can undergo bidirectional sex change, allowing them to overcome such risks and ensure continued reproduction (*1, 2*). As mammals lack such a strategy, developing techniques to regulate sex is crucial for ensuring mammalian reproduction and preserving biodiversity.

While cloning technology is an important tool for conserving endangered species (*3, 4*), its ability to facilitate or resume reproduction is constrained by the sex of resulting clones, as it typically produces offspring of the same sex as the donor cells. This poses a significant challenge, especially when the available genetic resources are limited in both quantity and diversity, as is often the case for extinct or endangered species (*5, 6*). Interestingly, a previous study suggested the possibility of resuming sexual reproduction through male somatic cell nuclear transfer, based on the finding that one female XO cloned mouse was accidentally born among 27 Sertoli cell-derived clones (*7*). However, no further progress has been made since this observation.

In mammals, sex is determined by the widely conserved Y-chromosome-linked gene Sry. Sry expression in the developing gonad triggers the activation of the male differentiation pathway and testis formation. In the absence of Sry, male-to-female sex reversal occurs in XY mouse embryos, resulting in the development of ovaries. However, the resulting XY ovaries often exhibit varying degrees of structural and functional abnormalities, and fertility is frequently compromised (*8-12*). By contrast, XO female mice are healthy and fertile (*13*). Thus, conversion from males to females through Y chromosome elimination in XY mouse embryos represents a promising strategy for sex reversal while securing female fertility. However, a robust method to induce Y chromosome elimination *in vivo*, as well as its application for mammalian reproduction, has not yet been established.

### Y chromosome elimination in mouse embryonic stem cells

We set out to establish a strategy to efficiently eliminate the Y chromosome *in vivo* to achieve robust sex reversal in mice. We reasoned that targeted inactivation of the Y chromosome centromere (Y-cent) in fertilized eggs would selectively eliminate Y chromosome in descendant cells, because loss of centromere function is not expected to activate the spindle assembly checkpoint, which monitors kinetochore-microtubule attachment (**Fig. 1A**). Indeed, previous studies have demonstrated successful Y chromosome removal in cultured cells either by using a mutant form of CENP-A (*14*) or by introducing CRISPR-Cas9-mediated double-strand breaks at the Y-cent (*15-18*). However, whether such chromosome-scale manipulation can be effectively applied in the context of animal reproduction remains unknown. To establish an efficient centromere-targeting strategy based on Y-cent cleavage, we designed three independent sgRNAs (Y1, Y2, and Y3) showing different numbers of target sites and efficiency scores (*19*) (**Tables S1 and S2**) and expressed them together with Cas9 in male mouse embryonic stem cells (mESCs). Genotyping PCR of individual colonies revealed that Y3 sgRNA exhibited the highest efficiency in eliminating the Y chromosome (**Fig. S1, A and B**). Combining all three sgRNAs did not improve efficiency, and therefore the single Y3 sgRNA was selected for subsequent experiments. To confirm whether the entire Y chromosome was eliminated, we performed chromosome-painting DNA-FISH (**Fig. 1B**). Y-chromosome painting signals were undetectable in a large population of cells transduced with Y3 sgRNA and Cas9, suggesting successful elimination of the entire chromosome (**Fig. 1, B and C**). For convenience, we hereafter refer to this Y chromosome elimination approach as Y-CUT (Y chromosome elimination via centromere-targeted Cas9-induced cuts).

**Figure 1.**
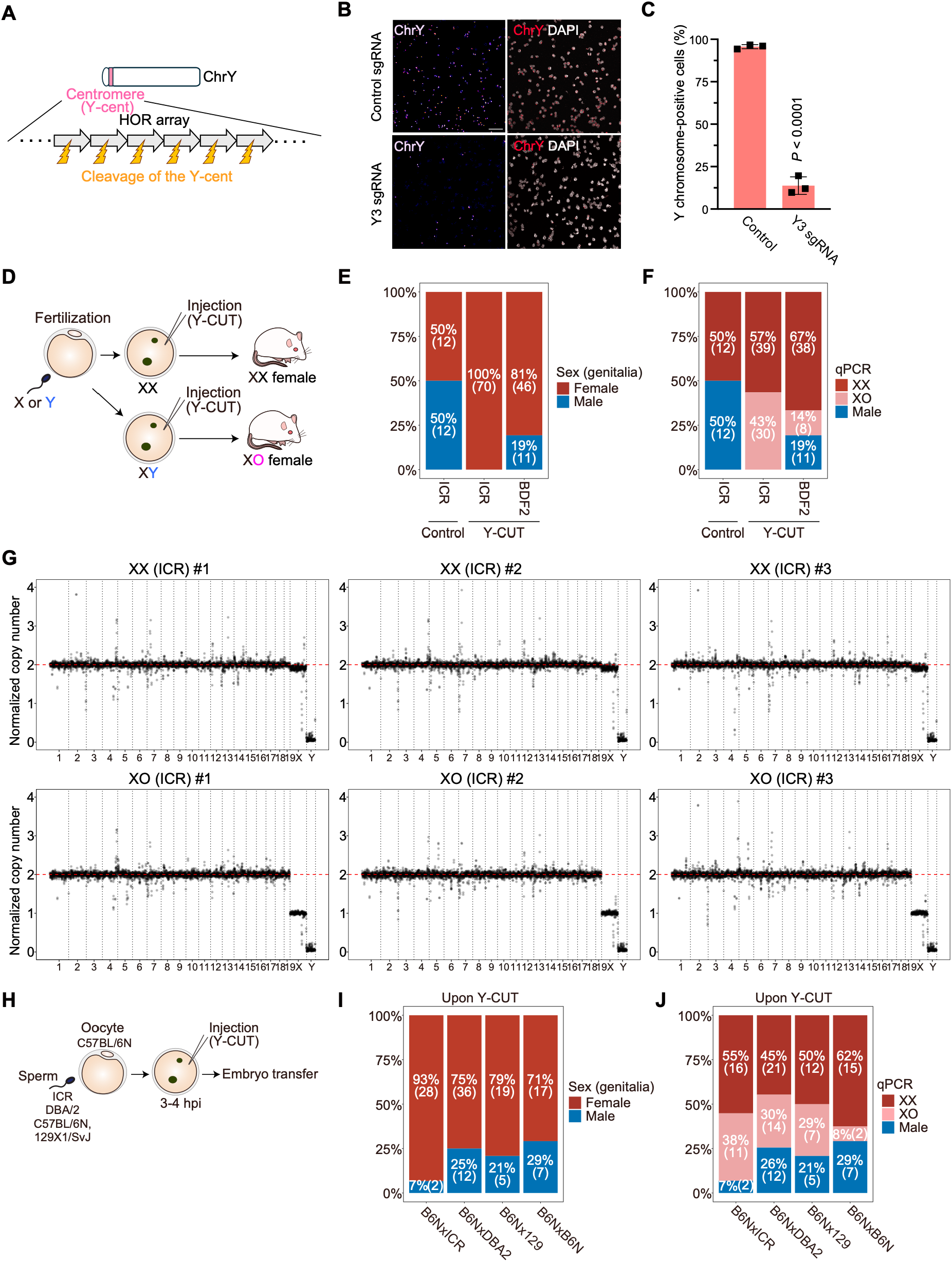
Targeted elimination of the Y chromosome in cultured cells and mice. **(A)** Schematic illustration of the Y-CUT method. HOR, higher-order repeat. **(B)** DNA-FISH for the Y chromosome in mESCs after the exogenous expression of control (GFP) or Y3 sgRNA together with Cas9. DAPI was used to stain DNA. Scale bars, 50 µm. **(C)** Bar plots showing the percentage of Y chromosome-positive mESCs. Results from three independent assays are shown. P values were calculated using a two-sided Mann–Whitney U test. **(D)** Schematic illustration showing the application of Y-CUT to mouse zygotes. **(E)** Bar plots showing the female and male ratio upon Y-CUT in mouse zygotes. Y-CUT was performed in ICR and BDF2 strains. Cas9 protein only was injected as a control. Results shown are pooled from at least two independent experiments. **(F)** Bar plots showing the XX and XO female and male ratio upon Y-CUT in mouse zygotes. Y-CUT was performed in ICR and BDF2 strains. Cas9 protein only was injected as a control. **(G)** Plots showing the chromosome copy numbers. XX or XO ICR mice were analyzed. Genome sequencing coverages at 1-Mb bins were plotted. Data was normalized to the XY genome sequencing data. **(H)** Experimental scheme for Y-CUT in different mouse genetic backgrounds. Sperm from different mouse strains were used, while oocytes were consistently collected from C57BL/6N female mice. **(I)** Bar plots showing the female and male ratio upon Y-CUT in mouse zygotes under indicated genetic backgrounds. Results shown are pooled from at least two independent experiments. **(J)** Bar plots showing the XX and XO ratio in females upon Y-CUT in mouse zygotes under indicated genetic backgrounds.

### Production of female-only offspring through Y-CUT

We next explored the applicability of Y-CUT for inducing sex reversal *in vivo*. To this end, we performed Y-CUT in mouse zygotes (**Fig. 1D**). Under the ICR genetic background, Y-CUT was remarkably efficient, consistently yielding female-only offspring, whereas control embryos (Cas9 only injection) produced males and females at comparable frequencies (**Fig. 1E**). As expected, nearly half (43%) of the females generated after Y-CUT exhibited an XO karyotype, as confirmed by qPCR, immunofluorescence of H3K27me3 (reflecting the absence of an inactive X chromosome in XO cells), and whole-genome sequencing (**Fig. 1, F and G; fig. S1, C and D**). Unexpectedly, Y-CUT efficiency was reduced in BDF2 embryos generated by intercrossing C57BL/6N × DBA/2 F1 (BDF1) mice, although Y chromosome elimination still occurred (**Fig. 1, E and F; fig. S1E**). This observation prompted us to investigate the underlying factors affecting Y-CUT efficiency.

Notably, two distinct types of Y chromosomes typically exist in laboratory mice. Although most inbred strains, including C57BL/6, DBA, 129, C3H, and BALB/c, possess a *Mus musculus (M*.*m) domesticus* nuclear genome, they carry a *M*.*m musculus*-type Y chromosome, which was introduced through East Asian “fancy” mice (*20*). In contrast, other *M*.*m. domesticus* strains, such as AKR and Swiss-derived lines (e.g., SJL, SWR, FVB, ICR, and CD1), harbor a *M*.*m. domesticus*-type Y chromosome. Consistent with this, we confirmed that ICR mice carry a *M*.*m. domesticus*-type Y chromosome, whereas other standard laboratory strains tested (C57BL/6J, C57BL/6N, C3H/He, BALB/c, DBA/2, and 129X1/SvJ) all carry a *M*.*m musculus*-type Y chromosome (**Fig. S1F**). To determine whether the *M*.*m. domesticus*-type Y chromosome is more susceptible to Y-CUT, we performed Y-CUT on zygotes generated by in vitro fertilization using C57BL/6N oocytes with sperm from C57BL/6N, DBA/2, 129X1/SvJ, or ICR males (**Fig. 1H**). As expected, Y chromosome elimination was markedly more efficient in zygotes from the C57BL/6N × ICR cross than in other combinations (**Fig. 1, I and J**). In cases of lower efficiency, partial XO females containing residual Y chromosome were observed, likely due to mosaic mutations (**Fig. S1G**). These differences in Y-CUT efficiency appear to reflect structural variation at the Y centromere: The *M*.*m. domesticus*-type Y centromere contains a 325–350 kb higher-order repeat (HOR) array composed of tandemly repeated 1.6 kb HOR units, whereas the *M*.*m. musculus*-type Y centromere consists of a shorter 90 kb HOR array with larger 2.3 kb HOR units (*20*). As a result, the *M. m. domesticus*-type Y centromere contains more than 400 predicted Y3 sgRNA target sites, with each 1.6-kb HOR unit containing two Y3 target sites (**Fig. S1H**). These findings identified the *M. m. domesticus*-type Y chromosome as the optimal substrate for robust Y-CUT, and all subsequent experiments were therefore performed using this genetic background.

### Y chromosome is released as micronuclei upon Y-CUT during early development

To investigate how the Y chromosome is eliminated upon Y-CUT *in vivo*, we performed DNA-FISH targeting the entire Y chromosome at the zygote, 2-cell, 4-cell, and morula stages (**Fig. 2A**). DNA-FISH for an X chromosome locus was conducted in parallel to distinguish embryos carrying one or two X chromosomes (**Fig. S2A**). At the zygote stage, Y chromosome signals were comparable to those in control embryos. However, at the 2-cell stage, the Y chromosome was either asymmetrically retained within the nucleus of a single blastomere or sequestered into micronuclei, with both phenotypes occurring at comparable frequencies (**Fig. 2, B and C; fig. S2, B and C; Movie S1**). At the 4-cell stage, Y chromosome micronuclei were detected in most embryos, although residual nuclear Y chromosome signals persisted in some blastomeres (**Fig. 2, B and C; fig. S2C; Movie S2**). At the morula stage, however, Y chromosome signals were absent from nuclei and were instead detected almost exclusively as micronuclei (**Fig. 2, B and C; Movie S3**). These Y chromosome micronuclei were not observed in control embryos (**Fig. S2D**). Across all developmental stages examined, the number of Y-chromosome–containing micronuclei per embryo was typically one or two, suggesting that Y-cent function was inactivated immediately after applying Y-CUT in zygotes and that the missegregated Y chromosome did not disperse into smaller fragments during development (**Fig. 2D**). Collectively, these observations indicate that the Y chromosome is ultimately released as micronuclei by the morula stage, likely due to stochastic missegregation during mitosis caused by centromere dysfunction (**Fig. S2E**). The resulting Y chromosome-containing micronuclei may subsequently undergo autophagic degradation (*21, 22*).

**Figure 2.**
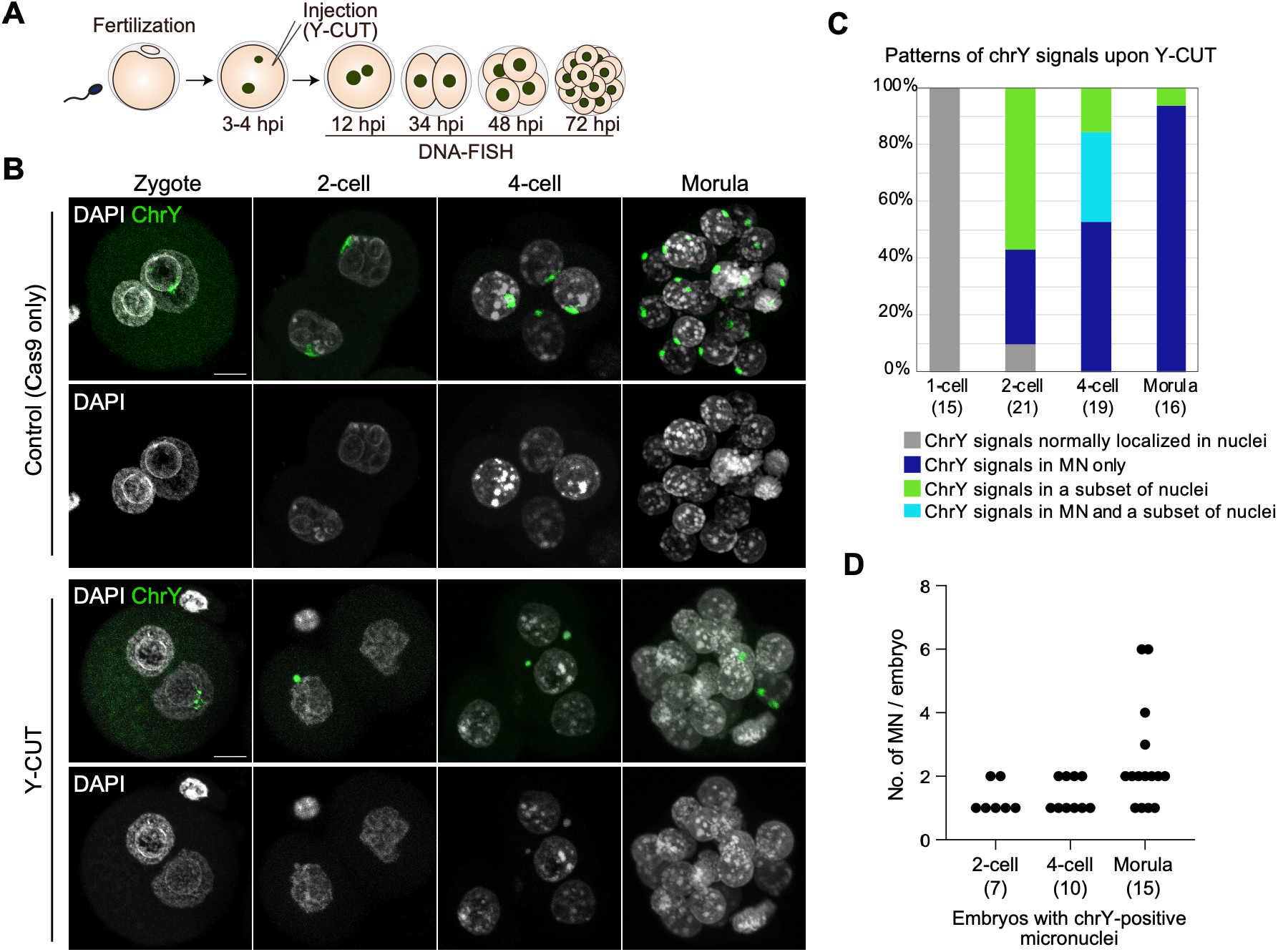
Y chromosome is released as micronuclei upon Y-CUT during early development. **(A)** Experimental scheme for DNA-FISH experiments after Y-CUT. **(B)** DNA-FISH for the Y chromosome in zygotes, 2-cell and 4-cell embryos, and morulae after Y-CUT. Cas9 protein only was injected in control. DAPI was used to stain DNA. Scale bars, 10 µm. See supplemental movies S1 to S3. **(C)** Stacked bar graph showing the frequency of the embryos showing the indicated Y chromosome signal localization patterns. The numbers of embryos analyzed are indicated. Results shown are pooled from at least three independent experiments. **(D)** Plots showing the numbers of Y chromosome-positive micronuclei (MN) per embryo upon Y-CUT in zygotes. Embryos carrying Y chromosome-positive micronuclei were selected for this analysis.

### Establishment of dual-sex cloning

Animal cloning through somatic cell nuclear transfer (SCNT) enables the generation of live organisms carrying the genome of the donor cell (*3*). However, SCNT typically produces cloned offspring of the same sex as the donor cells, thereby restricting the ability to resume reproduction from scarce genetic resources. Therefore, we sought to develop a strategy to generate both sexes using only XY somatic donor cells. Sertoli cells from neonatal male mice, which are commonly used as male donor cells (*23*), were initially used for SCNT. After nuclear transfer, Y-CUT was conducted by electroporation to avoid damage from repeated injections (**Fig. 3A**). In addition, a previously optimized protocol using HDAC and G9a inhibitors was integrated to maximize the SCNT efficiency (*24*). Using this approach, we efficiently obtained Y-CUT-applied 13 SCNT pups (11% birth rate per transferred embryos), and strikingly, ten of these (76.9%) were female, demonstrating the ability to produce female clones from male somatic donor cells (**Fig. 3B and fig. S3A**). All females were confirmed to be XO (**Fig. 3C and fig. S3B**), and both the female and male SCNT pups grew to adulthood (**Fig. 3D**). Furthermore, intercrossing these female and male clones produced the next generation (**Fig. 3, E and F**). The female offspring in this generation included karyotypically normal XX females, demonstrating restoration of the normal XX karyotype through sexual reproduction (**Fig. 3G and fig. S3C**). Slightly higher birth rate of XX females compared to XO females was consistent with the previous observation (*25, 26*) (**Fig. 3G**). Altogether, these results demonstrate that dual-sex cloning followed by reproduction can be achieved using only male somatic donor cells.

**Figure 3.**
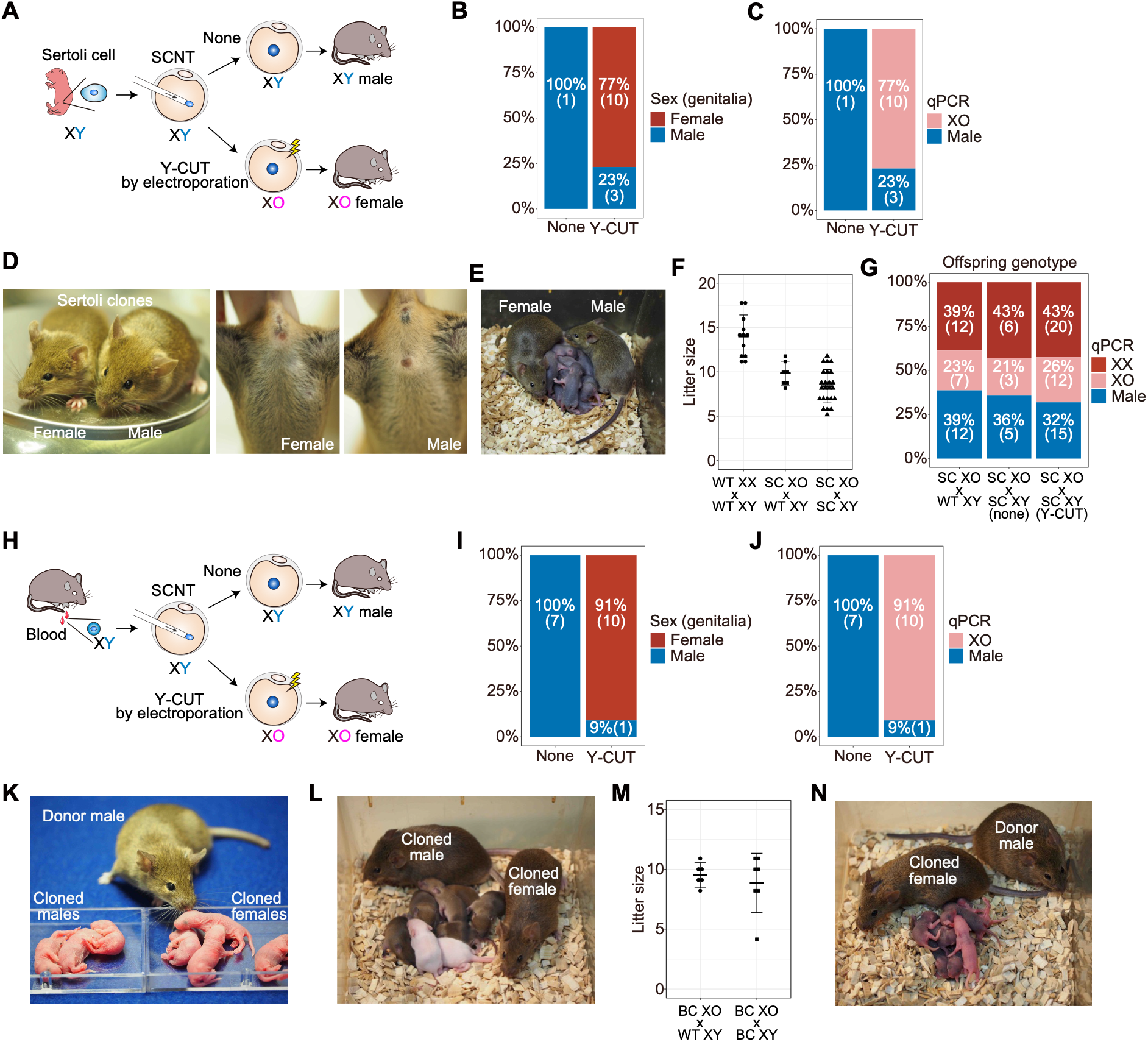
Parallel production of female and male mice from a male somatic genome by dual-sex cloning. **(A)** Experimental scheme for Y-CUT in somatic cell nuclear transfer. Sertoli (XY) cells from neonatal mice were used as donors. Y-CUT was performed by electroporation to avoid repeated injection. **(B)** Bar plots showing the female and male pup ratio upon Y-CUT after Sertoli cell nuclear transfer. Sex ratio of pups without Y-CUT (indicated as “none”) is also indicated as a control. **(C)** Bar plots showing the generation of XO females upon Y-CUT after Sertoli cell nuclear transfer. **(D)** Images showing the growth of female and male cloned mice to adulthood after Sertoli cell nuclear transfer. **(E)** Images showing the birth of next generation upon crosses between Sertoli cell nuclear transfer-derived female and male adult clones. **(F)** Plots showing the litter size upon crosses of indicated mice. WT XX, wild-type XX females; WT XY, wild-type XY males; SC XO, Sertoli cell nuclear transfer-derived XO females generated through Y-CUT; SC XY, Sertoli cell nuclear transfer-derived XY males. **(G)** Bar plots showing the XX and XO female and male ratio in offspring generated by crosses of indicated mice. **(H)** Experimental scheme for Y-CUT in somatic cell nuclear transfer. Male adult blood cells were used as donors. **(I)** Bar plots showing the female and male pup ratio upon Y-CUT after male blood cell nuclear transfer. Sex ratio of pups without Y-CUT (indicated as “none”) is also indicated as a control. **(J)** Bar plots showing the generation of XO females upon Y-CUT after male blood cell nuclear transfer. **(K)** Images showing the birth of male and female cloned pups generated using blood cells from the indicated adult male donor mouse. **(L)** Images showing the birth of next generation upon crossing male and female cloned mice produced by male blood cell nuclear transfer with or without Y-CUT. **(M)** Plots showing the litter size upon crosses of indicated mice. WT XY, wild-type XY males; BC XO, blood cell nuclear transfer-derived XO females generated through Y-CUT; BC XY, blood cell nuclear transfer-derived XY males. **(N)** Images showing the birth of next generation by crossing donor male and female cloned mice produced by male blood cell nuclear transfer with Y-CUT. The female cloned mouse was derived from a blood cell from the indicated donor male.

### Reproduction from a single adult male mouse genome

Having established the feasibility of dual-sex cloning, we next addressed whether reproduction could be achieved using adult male somatic cells while keeping the donor animals alive. To this end, we collected blood cells from the tail of a male adult mouse and used them as donor cells (*27*) (**Fig. 3H**). As expected, control SCNT without Y-CUT generated male pups only (seven male pups with 8% birth rate per transferred embryos). In contrast, ten females and one male were obtained after applying Y-CUT (8% birth rate per transferred embryos) (**Fig. 3, I and J; fig. S3, A and B**). This resulted in cloned males and females as well as the donor male all remaining alive (**Fig. 3K**). Intercrossing the cloned females and males gave rise to the next generation (**Fig. 3, L and M**). Remarkably, the original donor male successfully initiated reproduction by mating with female clones derived from his own somatic genome (**Fig. 3N**). Similar results were obtained by using tail tip fibroblast as donor cells (**Fig. S3, A and B**). These results demonstrate that the dual-sex cloning approach enables reproduction from a single adult male.

### Reproduction from cryopreserved male somatic cells

Dual-sex cloning offers a promising strategy for restoring reproduction using male somatic cells as genetic resources. However, somatic cells are not always obtainable from live animals. In practice, cells from deceased individuals or endangered species, are often cryopreserved. To determine whether reproduction can be resumed from freeze-stored male somatic cells, we performed dual-sex cloning using cryopreserved blood cells derived from adult male mice (**Fig. 4A**). Using male blood cells that had been frozen for 1 day or 1 month, control SCNT produced only male pups (six and three male pups, corresponding to 14% and 5% birth rates per transferred embryos, respectively), whereas Y-CUT–treated SCNT generated only female pups (two and five female pups, corresponding to 5.7% and 3.9% birth rates, respectively) (**Fig. 4B; Fig. S3, A and B**). Importantly, these female and male clones produced the next generation upon intercrossing (**Fig. 4C**). These results demonstrate that dual-sex cloning enables reproduction from cryopreserved male somatic cells, highlighting its utility as a practical strategy for restoring reproduction.

**Figure 4.**
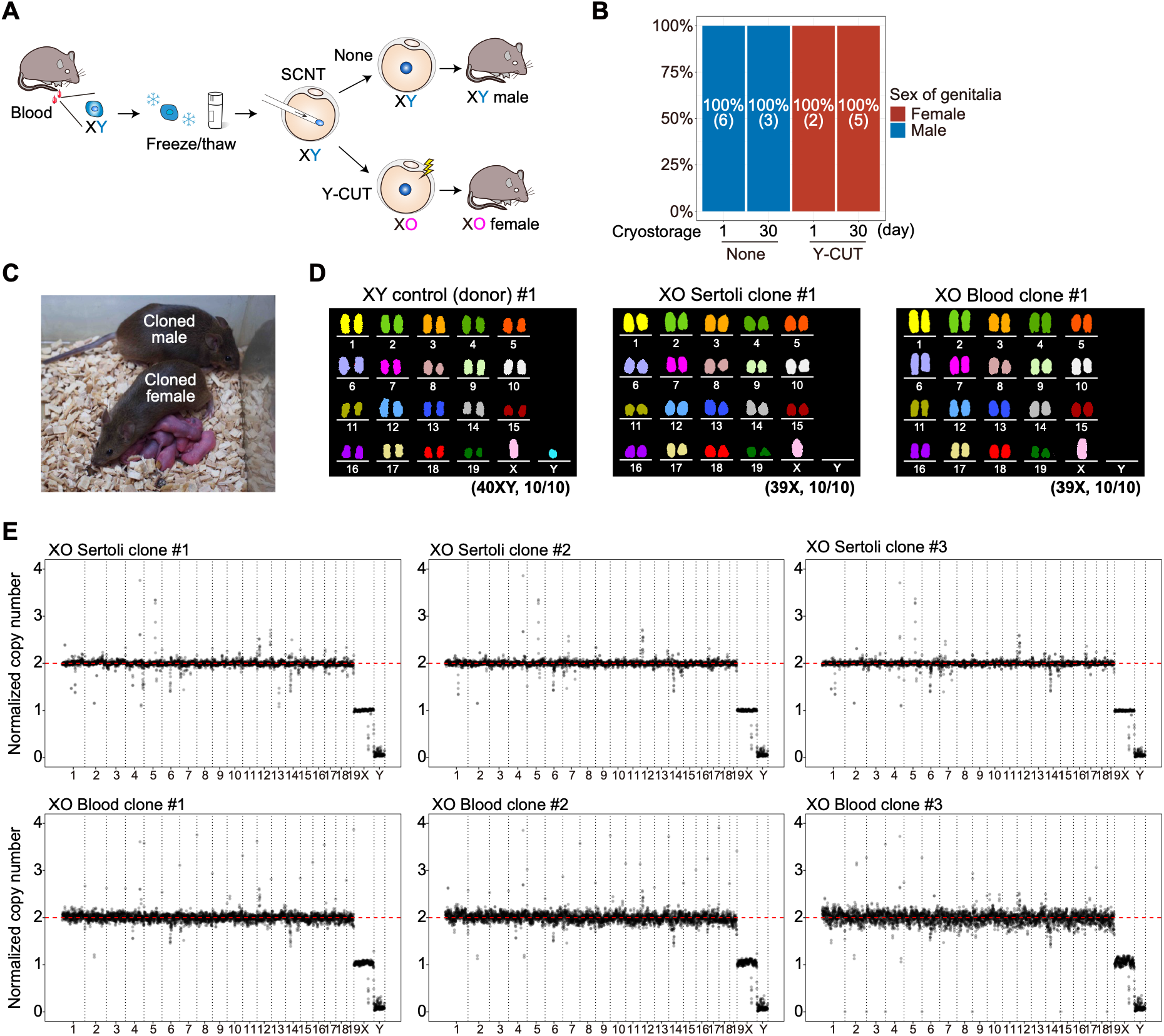
Initiation of reproduction from frozen male somatic cells. **(A)** Experimental scheme for Y-CUT in somatic cell nuclear transfer. Freeze-stored adult male blood cells or tail-tip fibroblasts mice were used as donors. Y-CUT was performed by electroporation to avoid repeated injection. **(B)** Bar plots showing the female and male pup ratio upon Y-CUT after nuclear transfer of freeze-stored adult male blood cells. The donor cells were freeze-stored for 1 day or 30 days. Sex ratio of pups without Y-CUT (indicated as “none”) is also indicated as a control. **(C)** Images showing the birth of next generation upon crosses between freeze-stored male blood cell nuclear transfer-derived female and male adult clones. **(D)** Karyotypes of male donor mice and cloned female mice generated through Y-CUT. Multicolor FISH was performed to examine their karyotype. All ten examined somatic cells in donor mice showed normal karyotype (40XY), while those in Sertoli cell- or male blood cell-derived cloned female mice generated through Y-CUT indicated XO (39X) karyotype. No Robertsonian or reciprocal translocations were observed except for the complete loss of the Y chromosome in the cloned female mice. **(E)** Plots showing the chromosome copy numbers in cloned female mice generated through Y-CUT. Genome sequencing coverage at 1-Mb bins was plotted. Data was normalized to the XY genome sequencing data. Three independent Sertoli cell- or male blood cell-derived cloned female mice were analyzed.

### Genomic stability of sex-reversed female clones generated by dual-sex cloning

Dual-sex cloning involves both centromere-targeted chromosome cleavage and the SCNT procedure, raising concerns about the unintended induction of structural chromosomal abnormalities, such as Robertsonian or reciprocal translocations, as well as off-target mutations. Because balanced chromosomal rearrangements are compatible with viability and fertility, such alterations could be transmitted to subsequent generations. To evaluate these potential risks, we first performed multicolor FISH for detailed karyotypic analysis. Examination of three control XY donor males confirmed normal karyotypes. Analysis of three Sertoli cell-derived and three blood-derived XO female clones revealed specific elimination of the Y chromosome without Robertsonian or reciprocal translocations (**Fig. 4D and fig. S4A**). All metaphases examined exhibited uniform karyotypes, with no indication of mosaicism. To complement these cytogenetic analyses, we next performed whole-genome sequencing at 8–11× coverage. Comparative analyses between XY controls and XO female clones confirmed normal copy numbers of all autosomes and the X chromosome, together with complete loss of the Y chromosome in Y-CUT-treated XO females (**Fig. 4E**). Analysis of predicted off-target sites revealed sequence alterations at fewer than 1% of the examined loci (**Fig. S4, B and C**). All detected off-target events were mosaic mutations consisting of 1–10 bp indels located outside exonic regions. Collectively, these findings demonstrate that dual-sex cloning does not introduce detectable large-scale genomic abnormalities while enabling the parallel generation of male and female individuals from a single male somatic genome.

### A programmable platform for Y chromosome elimination and sex reversal

Based on the high efficiency of Y-CUT, we finally explored whether this technology could be genetically encoded to establish a programmable platform for Y chromosome elimination *in vivo*. To this end, we generated knock-in (KI) mice carrying a ROSA26-targeted cassette expressing the Y3 sgRNA together with Cre-dependent Cas9 (**Fig. 5A and fig. S5A**). Heterozygous KI/WT mESCs derived from these mice efficiently eliminated the Y chromosome following transient expression of Cre recombinase, confirming the functionality of the integrated cassette (**Fig. S5B**). We next asked whether genetically encoded Y-CUT could be temporally controlled. KI/WT mESCs stably expressing MerCreMer, a tamoxifen-inducible Cre recombinase, were treated with 4-hydroxytamoxifen (4-OHT) (**Fig. 5B**). Y-chromosome–containing micronuclei were frequently observed 24 h after 4-OHT treatment, whereas Y chromosome signals were almost completely lost by 72–96 h (**Fig. 5, C and D**), demonstrating temporal control of Y chromosome elimination. Finally, we tested whether genetically encoded Y-CUT could induce Y chromosome elimination *in vivo* by crossing homozygous KI/KI mice with Sox2-Cre/+ mice, which express Cre throughout the embryonic lineages (*28*) (**Fig. 5E**). As anticipated, the offspring were female-biased (**Fig. 5F**), and karyotype analysis revealed that the half of the Sox2-Cre–positive females exhibited an XO karyotype (**Fig. 5, G and H**). Collectively, these findings establish genetically encoded Y-CUT as a programmable platform for conditional chromosome elimination and male-to-female sex reversal.

**Figure 5.**
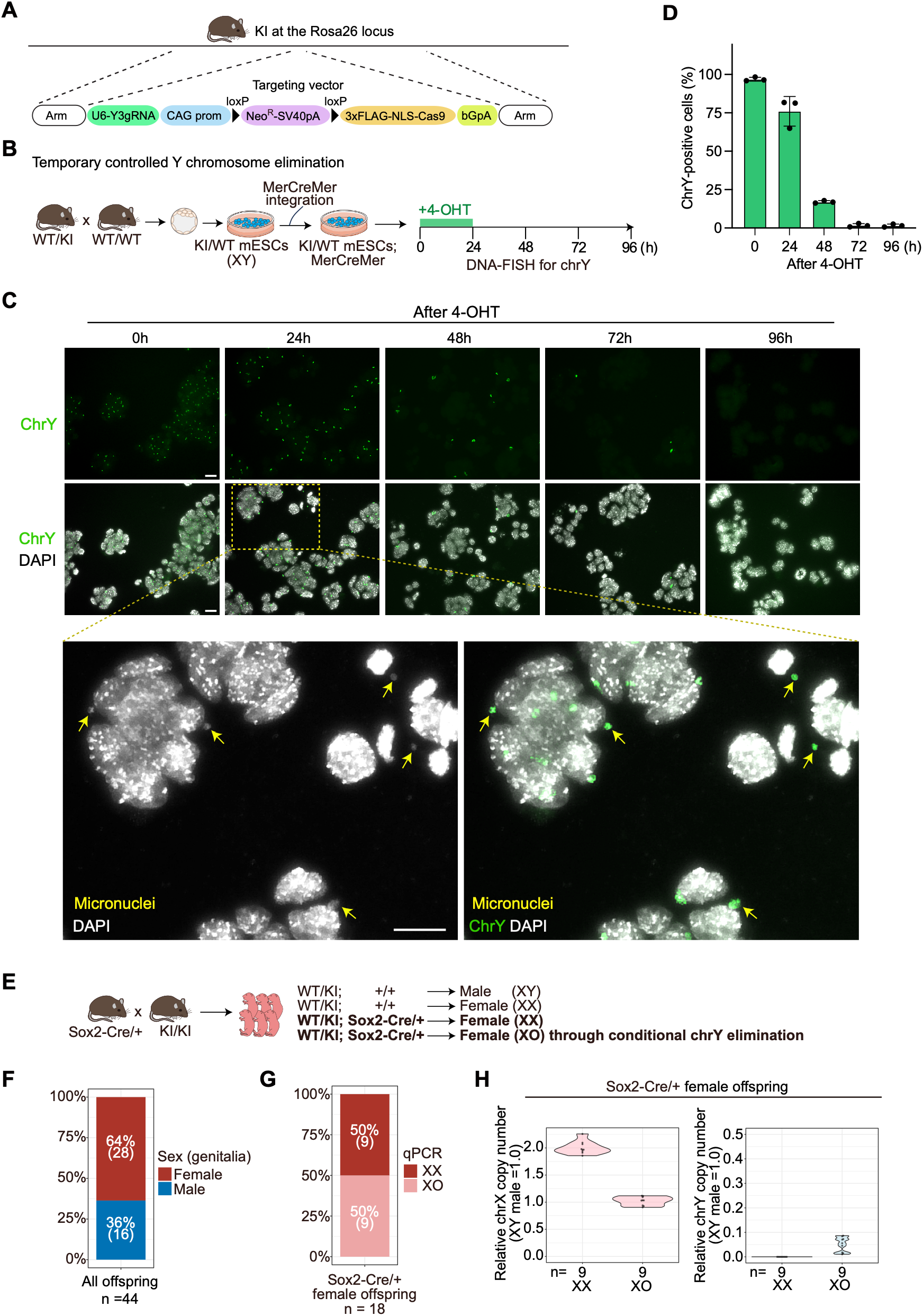
Genetic encoding of Y-CUT enables conditional chromosome removal and male-to-female sex reversal. **(A)** Knock-in of a genetically encoded Y-CUT cassette at the ROSA26 locus in mice. Y3 sgRNA is constitutively expressed under a U6 promoter, while Cas9 expression is suppressed by a floxed neomycin-resistance cassette before Cre-mediated recombination. Cre-mediated recombination excises the neomycin cassette and allows Cas9 expression. **(B)** Experimental scheme for a temporally controlled Y chromosome elimination in mESCs. Heterozygous knock-in (KI/WT) mESCs carrying the genetically encoded Y-CUT cassette was derived from blastocysts generated by the cross between heterozygous KI mice and wild-type mice. KI/WT mESCs expressing MerCreMer were then generated. **(C)** DNA-FISH for the Y chromosome was performed at 0 (without 4-OHT), 24, 48, 72, and 96 h after the 4-OHT treatment for 24 h. The area indicated by a dashed square is magnified below. Y chromosome-positive micronuclei (indicated by yellow arrows) were already observed 24 h after the 4-OHT treatment. Scale bars, 20 µm. **(D)** Bar plots showing the percentage of Y-chromosome positive cells after 4-OHT treatment. Results shown are from three independent experiments. **(E)** Experimental scheme for conditional Y chromosome elimination. Sox2-Cre/+ mice were crossed with homozygous knock-in mice. High female pup ratio is expected as XY offspring carrying the Sox2-Cre allele are expected to become XO females. **(F)** Bar plots showing the female and male pup ratio upon crosses indicated in (E). **(G)** Bar plots showing the XX and XO female ratio in Sox2-Cre/+ female offspring generated by crosses indicated in (E). **(H)** Violin plots showing copy numbers of the X chromosome (left) and Y chromosome (right) in XX and XO female offspring carrying Sox2-Cre. Chromosome copy numbers were determined by qPCR.

## Discussion

Here we establish dual-sex cloning, a strategy that enables the generation of both male and female individuals from a single male somatic genome. By combining efficient Y chromosome elimination with optimized somatic cell nuclear transfer, we generated fertile XO female clones alongside XY male clones from the same male genome. These animals produced offspring through mating between male and female clones. Moreover, an adult male donor produced offspring by mating with a female clone derived from its own somatic cells. The resulting progeny included chromosomally normal XX females, demonstrating that a conventional XX–XY reproductive cycle can be restored in the next generation. Unlike approaches that alter sex ratios by inducing sex-specific embryonic lethality (*29, 30*), dual-sex cloning changes the sexual identity of viable cloned animals through targeted chromosome elimination. These findings show that the sex of a somatic-cell donor is no longer an absolute constraint on the sex of its cloned descendants and establish a new route for initiating mammalian sexual reproduction from a single male genome.

Dual-sex cloning may have practical applications in both the generation of genetically engineered animal models and the preservation of limited genetic resources (*3*). In conventional SCNT-based workflows, genome editing can be completed and validated in donor cells before nuclear transfer, but the resulting founders are restricted to the sex of the donor cells, often requiring additional generations of breeding before both sexes carrying the desired genotype become available. By enabling genetically matched male and female founders to be generated directly from a single engineered donor cell, dual-sex cloning could substantially accelerate colony establishment, reduce breeding efforts, and facilitate the establishment of complex genetically engineered models requiring multiple engineered alleles or sequential genome modifications. The benefits of this strategy are expected to be particularly pronounced in species with long generation intervals, including non-human primates, where each breeding generation requires several years (*31, 32*). Beyond experimental animal production, dual-sex cloning may also expand the reproductive potential of cryopreserved somatic cells. Our findings demonstrate that fertile male and female clones can be generated from frozen male cells and subsequently used to produce offspring, providing a potential strategy for reconstructing reproduction when only male somatic material remains available. Such an approach may complement existing biobanking and conservation efforts (*33*) by maximizing the reproductive utility of limited genetic resources. Taken together, these features make dual-sex cloning particularly attractive for species in which conventional breeding is slow, expensive, or impractical.

Several barriers must nevertheless be overcome before dual-sex cloning can be extended to a broader range of mammals. Although XO females are fertile in mice and potentially in other rodents, they are infertile in other mammalian species, including humans and horses (*34, 35*). This species difference has been attributed to variation in the structure and gene content of the sex chromosomes (*34*). Thus, additional technological advances will be required to extend dual-sex cloning-mediated reproduction to a broader range of species. Furthermore, dual-sex cloning currently relies on enucleated oocytes for nuclear transfer, necessitating the use of female-derived cellular materials. A possible strategy for initiating reproduction solely from male-derived materials may involve generating oocytes from XY male pluripotent stem cells (*36*). Integrating such developmental engineering approaches with dual-sex cloning will further expand the possibilities for mammalian reproduction.

Beyond enabling dual-sex cloning, Y-CUT also establishes a general framework for programmable chromosome engineering *in vivo*. Whereas recent advances in genome editing have transformed our ability to manipulate DNA sequences with high efficiency and precision (*37*), technologies capable of chromosome-scale manipulation, particularly *in vivo*, remain scarce. By targeting centromere function, genetically encoded Y-CUT enables controlled elimination of a whole chromosome while retaining compatibility with spatiotemporal regulation. This capability provides a versatile experimental system for investigating Y chromosome biology, including mosaic loss of the Y chromosome and Y chromosome dosage effects (*38, 39*).

Collectively, dual-sex cloning expands the conceptual boundaries of mammalian reproduction by demonstrating that sexual reproduction can be re-initiated from a single male genome. Beyond its immediate applications in reproductive biology and conservation, this work also establishes the Y-CUT platform as a general framework for programmable chromosome elimination and future chromosome engineering.

## Acknowledgments

We would like to thank the members of our laboratories for discussion, Narumi Ogonuki and Michiko Hirose for technical support, Yusuke Miyanari for technical advice, Masahito Ikawa for providing the EGR-G01 mESC cell line, and Hitoshi Niwa for providing a MerCreMer expression plasmid. We also thank the animal facility for animal care and for providing access to shared microscopy equipment.

## Funding

This work was supported by grants from MEXT Grant-in-Aid for Transformative Research Areas (JP25H01354 to T.I.; JP25H01356 to S.M.), Uehara Memorial Foundation (T.I.), JST FOREST Program (JPMJFR243B to T.I.; JPMJFR221G to S.M.), JSPS KAKENHI Grant-in-Aid for Scientific Research (B) (JP24K01947 to T.I.; JP25K02201 to S.M.), and JSPS KAKENHI Grant-in-Aid for Scientific Research (C) (JP25K09494 to T.H.).

## Author contributions

Conceptualization: TI, SM

Methodology: TI, SM

Investigation: SM, NU, MS, SK, DQP, A.Okamoto, AI, TA, HS, KI, TH, TI

Visualization: SM, NU, MS, SK, DQP, A.Okamoto, TH, TI

Funding acquisition: TI, SM, TH

Project administration: TI, SM

Supervision: TI, SM, KI, A.Ogura

Writing – original draft: TI, SM

Writing – review & editing: TI, SM, TH, KI, A.Ogura

## Competing interests

The authors declare a patent application related to this work.

## Data, code, and materials availability

The sequencing data from this study are available at the SRA, accession number PRJNA1498686.

## Supplementary Materials

Materials and Methods

Figs. S1 to S5

Tables S1 to S2

Movies S1 to S3

## References

1. B. Capel, Vertebrate sex determination: evolutionary plasticity of a fundamental switch. Nat Rev Genet 18, 675–689 (2017).

2. Y. Nagahama, T. Chakraborty, B. Paul-Prasanth, K. Ohta, M. Nakamura, Sex determination, gonadal sex differentiation, and plasticity in vertebrate species. Physiol Rev 101, 1237–1308 (2021).

3. S. Matoba, Y. Zhang, Somatic Cell Nuclear Transfer Reprogramming: Mechanisms and Applications. Cell Stem Cell 23, 471–485 (2018).

4. A. Ogura, K. Inoue, T. Wakayama, Recent advancements in cloning by somatic cell nuclear transfer. Philos Trans R Soc Lond B Biol Sci 368, 20110329 (2013).

5. A. Mooney et al., Maximizing the potential for living cell banks to contribute to global conservation priorities. Zoo Biol 42, 697–708 (2023).

6. K. Theissinger et al., How genomics can help biodiversity conservation. Trends Genet 39, 545–559 (2023).

7. K. Inoue et al., Sex-reversed somatic cell cloning in the mouse. J Reprod Dev 55, 566–569 (2009).

8. J. Gubbay et al., A gene mapping to the sex-determining region of the mouse Y chromosome is a member of a novel family of embryonically expressed genes. Nature 346, 245–250 (1990).

9. R. Lovell-Badge, E. Robertson, XY female mice resulting from a heritable mutation in the primary testis-determining gene, Tdy. Development 109, 635–646 (1990).

10. T. Kato et al., Production of Sry knockout mouse using TALEN via oocyte injection. Sci Rep 3, 3136 (2013).

11. H. Wang et al., TALEN-mediated editing of the mouse Y chromosome. Nat Biotechnol 31, 530–532 (2013).

12. A. Sakashita et al., XY oocytes of sex-reversed females with a Sry mutation deviate from the normal developmental process beyond the mitotic stagedagger. Biol Reprod 100, 697–710 (2019).

13. B. M. Cattanach, XO mice. Genetics Research 3, 487–490 (1962).

14. P. Ly et al., Selective Y centromere inactivation triggers chromosome shattering in micronuclei and repair by non-homologous end joining. Nat Cell Biol 19, 68–75 (2017).

15. F. Adikusuma, N. Williams, F. Grutzner, J. Hughes, P. Thomas, Targeted Deletion of an Entire Chromosome Using CRISPR/Cas9. Mol Ther 25, 1736–1738 (2017).

16. T. A. Prowse, F. Adikusuma, P. Cassey, P. Thomas, J. V. Ross, A Y-chromosome shredding gene drive for controlling pest vertebrate populations. Elife 8, (2019).

17. S. Sano et al., Hematopoietic loss of Y chromosome leads to cardiac fibrosis and heart failure mortality. Science 377, 292–297 (2022).

18. H. A. Abdel-Hafiz et al., Y chromosome loss in cancer drives growth by evasion of adaptive immunity. Nature 619, 624–631 (2023).

19. J. G. Doench et al., Rational design of highly active sgRNAs for CRISPR-Cas9-mediated gene inactivation. Nat Biotechnol 32, 1262–1267 (2014).

20. M. D. Pertile, A. N. Graham, K. H. Choo, P. Kalitsis, Rapid evolution of mouse Y centromere repeat DNA belies recent sequence stability. Genome Res 19, 2202–2213 (2009).

21. S. Rello-Varona et al., Autophagic removal of micronuclei. Cell Cycle 11, 170–176 (2012).

22. K. Krupina, A. Goginashvili, D. W. Cleveland, Causes and consequences of micronuclei. Curr Opin Cell Biol 70, 91–99 (2021).

23. A. Ogura et al., Production of male cloned mice from fresh, cultured, and cryopreserved immature Sertoli cells. Biol Reprod 62, 1579–1584 (2000).

24. S. Matoba et al., Reduction of H3K9 methylation by G9a inhibitors improves the development of mouse SCNT embryos. Stem Cell Reports 19, 906–921 (2024).

25. R. LeMaire-Adkins, P. A. Hunt, Nonrandom segregation of the mouse univalent X chromosome: evidence of spindle-mediated meiotic drive. Genetics 156, 775–783 (2000).

26. F. E. Clark, T. Akera, Unravelling the mystery of female meiotic drive: where we are. Open Biol 11, 210074 (2021).

27. S. Kamimura et al., Mouse cloning using a drop of peripheral blood. Biol Reprod 89, 24 (2013).

28. S. Hayashi, P. Lewis, L. Pevny, A. P. McMahon, Efficient gene modulation in mouse epiblast using a 8 Sox2Cre transgenic mouse strain. Mech Dev 119 Suppl 1, S97–S101 (2002).

29. I. Yosef et al., A genetic system for biasing the sex ratio in mice. EMBO Rep 20, e48269 (2019).

30. C. Douglas et al., CRISPR-Cas9 effectors facilitate generation of single-sex litters and sex-specific phenotypes. Nat Commun 12, 6926 (2021).

31. E. Sasaki, Prospects for genetically modified non-human primate models, including the common marmoset. Neurosci Res 93, 110–115 (2015).

32. J. K. Schmidt, K. M. Jones, T. Van Vleck, M. E. Emborg, Modeling genetic diseases in nonhuman primates through embryonic and germline modification: Considerations and challenges. Sci Transl Med 14, eabf4879 (2022).

33. O. A. Ryder, M. Onuma, Viable Cell Culture Banking for Biodiversity Characterization and Conservation. Annu Rev Anim Biosci 6, 83–98 (2018).

34. T. Raudsepp, P. J. Das, F. Avila, B. P. Chowdhary, The pseudoautosomal region and sex chromosome aneuploidies in domestic species. Sex Dev 6, 72–83 (2012).

35. C. H. Gravholt et al., Clinical practice guidelines for the care of girls and women with Turner syndrome. Eur J Endocrinol 190, G53–G151 (2024).

36. K. Murakami et al., Generation of functional oocytes from male mice in vitro. Nature 615, 900–906 (2023).

37. M. Pacesa, O. Pelea, M. Jinek, Past, present, and future of CRISPR genome editing technologies. Cell 187, 1076–1100 (2024).

38. A. S. F. Berry et al., A genome-first study of sex chromosome aneuploidies provides evidence of Y chromosome dosage effects on autism risk. Nat Commun 15, 8897 (2024).

39. B. Bruhn-Olszewska et al., The effects of loss of Y chromosome on male health. Nat Rev Genet 26, 320–335 (2025).

40. Y. Fujihara, K. Kaseda, N. Inoue, M. Ikawa, M. Okabe, Production of mouse pups from germline transmission-failed knockout chimeras. Transgenic Res 22, 195–200 (2013).

41. S. Bae, J. Park, J. S. Kim, Cas-OFFinder: a fast and versatile algorithm that searches for potential off-target sites of Cas9 RNA-guided endonucleases. Bioinformatics 30, 1473–1475 (2014).

42. S. Fujii et al., Nr0b1 is a negative regulator of Zscan4c in mouse embryonic stem cells. Sci Rep 5, 9146 (2015).

43. S. Matoba et al., RNAi-mediated knockdown of Xist can rescue the impaired postimplantation development of cloned mouse embryos. Proc Natl Acad Sci U S A 108, 20621–20626 (2011).

44. S. Matoba et al., Embryonic development following somatic cell nuclear transfer impeded by persisting histone methylation. Cell 159, 884–895 (2014).

45. N. Auer et al., ChromaWizard: An open source image analysis software for multicolor fluorescence in situ hybridization analysis. Cytometry A 93, 749–754 (2018).

46. C. D’Hulst, I. Parvanova, D. Tomoiaga, M. L. Sapar, P. Feinstein, Fast quantitative real-time PCR-based screening for common chromosomal aneuploidies in mouse embryonic stem cells. Stem Cell Reports 1, 350–359 (2013).

47. G. Jun, M. K. Wing, G. R. Abecasis, H. M. Kang, An efficient and scalable analysis framework for variant extraction and refinement from population-scale DNA sequence data. Genome Res 25, 918–925 (2015).

48. A. McKenna et al., The Genome Analysis Toolkit: a MapReduce framework for analyzing next-generation DNA sequencing data. Genome Res 20, 1297–1303 (2010).

49. V. Boeva et al., Control-FREEC: a tool for assessing copy number and allelic content using next-generation sequencing data. Bioinformatics 28, 423–425 (2012).

50. B. S. Pedersen, A. R. Quinlan, Mosdepth: quick coverage calculation for genomes and exomes. Bioinformatics 34, 867–868 (2018).

51. A. R. Quinlan, I. M. Hall, BEDTools: a flexible suite of utilities for comparing genomic features. Bioinformatics 26, 841–842 (2010).

52. H. Li, A statistical framework for SNP calling, mutation discovery, association mapping and population genetical parameter estimation from sequencing data. Bioinformatics 27, 2987–2993 (2011).

53. J. T. Robinson et al., Integrative genomics viewer. Nat Biotechnol 29, 24–26 (2011).

