## Supplemental Materials for "Initiating mammalian reproduction from a single male genome"

**The PDF file includes:**

Materials and Methods
Figs. S1 to S5

**Other Supplementary Materials for this manuscript include the following:**

Tables S1 to S2
Movies S1 to S3

#### Materials and Methods

##### Animals

All animal experiments were approved by the Animal Experiments Committee of University of Yamanashi and RIKEN Tsukuba, and performed according to the guidelines for animal experiments at each institution. ICR, C57BL/6N (B6N), DBA/2, and B6N x DBA/2 F1 (BDF1) mice were purchased from Japan SLC Inc. 129/Sv mice were provided by RIKEN BRC through the National BioResource Project of MEXT/AMED, Japan (RBRC00002). Sox2-Cre mice are from the Jackson Laboratory (#008454). For embryo-transfer surgery, mice were anesthetized by intraperitoneal administration of medetomidine (0.3 mg $kg^{-1}$; Nippon Zenyaku Kogyo), midazolam (4 mg $kg^{-1}$; Maruishi Pharmaceutical), and butorphanol (5 mg $kg^{-1}$; Meiji Animal Health). Anesthesia was reversed by intraperitoneal administration of atipamezole (0.3 mg $kg^{-1}$; Nippon Zenyaku Kogyo) after surgery. Mice were housed in cages under specific pathogen-free conditions and had free access to water and food.

##### mESC culture

B6N x 129 F1 male mESCs (EGR-G01) (*40*) were cultured with medium consisting of DMEM (Nacalai tesque) containing 15% FBS (Thermo Fisher Scientific), 0.1 mM 2-mercaptoethanol (Thermo Fisher Scientific), nonessential amino acid (Thermo Fisher Scientific), sodium pyruvate (Thermo Fisher Scientific), penicillin/streptomycin (Thermo Fisher Scientific), leukemia inhibitory factor (Nacalai tesque), 0.4 μM PD0325901 (Fujifilm) and 3 μM CHIR99021 (Fujifilm) on gelatin-coated plates.

##### Y-CUT in mESC

425 gRNA sequences targeting the Y chromosome centromere were designed and selected based on target specificity, the numbers of target sites, AT content, and predicted efficiency scores (*19*). Their specificity was evaluated using Cas-OFFinder (*41*). Three independent gRNAs, each targeting different numbers of sites specifically within the Y centromere, were tested for their ability to eliminate the Y chromosome. gRNA target sequences were cloned into pX459 v2.0 (Addgene # 62988, a gift from Feng Zhang). These pX459 v2.0 plasmids were transfected into mESCs using lipofectamine 2000 (Thermo Fisher Scientific). 24 h after the transfection, cells were selected with 2 μg/mL puromycin for 24 h and then dissociated to form colonies from single cells. After colonies were formed in three to four days of culture, each colony was picked up and subjected to PCR genotyping. For DNA-FISH, cells cultured for 3 days after the puromycin selection were used. The following primers, which yield different size of PCR fragments from the X and Y chromosomes, were used.
F: TGGATGGTGTGGCCAATG
R: CACCTGCACGTTGCCCTT

##### Generation of knock-in mice

A targeting vector was constructed by inserting ROSA26 homology arms and CAG-loxP-Neo-pA-lox sequences to Y3 sgRNA expressing pX330 (Addgene # 42230, a gift from Feng Zhang). The targeting vector was used to generate knock-in mESCs. These knock-in mESCs were injected into 8-cell embryos to generate chimeric mice. Germline transmission was confirmed by PCR. For experiments, ICR males were crossed to target the *M.m. domesticus*-type Y chromosome.

**Generation of knock-in mESCs**

Blastocysts obtained from the crosses between heterozygous female KI mice and WT ICR male mice were used for mESC derivation. Blastocysts were placed to a gelatin-coated culture plates containing mESC culture medium described above, and primary ESC colonies were dissociated into single cells to establish mESC lines. Cre or MerCreMer-IRES-Hygro expression vectors (*42*) were transfected by lipofectamine 2000. 300 μg/mL hygromycin selection followed by single colony pickup was performed to generate MerCreMer stable transfectants.

**Determination of mouse Y chromosome origin**

The Y chromosome origin was determined by PCR as described previously (*20*). Phenol/chloroform-extracted genomic DNA from C57BL/6J, C57BL/6N, C3H/He, BALB/c, DBA/2, 129X1/SvJ, and ICR adult male mice was used as a template DNA. PCR was conducted using forward primer (5'-CACAGTGTAGAACACCGTAC-3') and reverse primer (5'-CGTTTCTCATTATATATGTTTTTCTTC-3'), which generated 0.82- and 1.6-kb PCR products from *M.m. musculus*-type and *M.m. domesticus*-type Y chromosome, respectively.

**In vitro fertilization (IVF)**

Spermatozoa were collected from the cauda epididymis of adult male mice and incubated in HTF drops for 1 h at 37°C under 5% CO2 in humidified air. Cumulus-oocyte-complexes (COCs) were collected from adult female mice by superovulation with injection of 5 IU pregnant mare serum gonadotropin (PMSG) followed by 7.5 IU human chorionic gonadotropin (hCG; Aska Pharmaceutical Co. Ltd.) at 48 h intervals. After 1 h of preincubation, the activated spermatozoa were added to the HTF drops containing COCs to initiate insemination. After 4–6 h of co-incubation, the fertilized zygotes were washed and cultured in drops of potassium-enriched simplex optimization medium (KSOM).

**Y-CUT in mouse zygotes**

Zygotes obtained by IVF were subjected to Y chromosome elimination at the pronuclear stage by introducing Cas9 ribonucleoprotein (RNP) complexes targeting the Y chromosome centromere. Y3 crRNA and tracrRNA (Alt-R CRISPR-Cas9 system, Integrated DNA Technologies) were annealed according to the manufacturer's instructions to generate a crRNA–tracrRNA duplex, which was subsequently mixed with recombinant Cas9 protein (Alt-R S.p. Cas9 Nuclease V3, Integrated DNA Technologies). Final concentrations were 100 ng/μL Cas9 and 50 ng/μL crRNA–tracrRNA. Cas9 RNPs were introduced into zygotes either by microinjection or by electroporation. For microinjection, approximately 5–10 pL of the RNP solution was injected to cytoplasm of zygotes using a piezo-assisted micromanipulator. For electroporation, zygotes were transferred into Opti-MEM (Thermo Fisher Scientific) containing Cas9 RNPs and electroporated using a NEPA21 electroporator (Nepa Gene, Chiba, Japan) equipped with a 1-mm gap electrode (CUY501P1-1.5, Nepa Gene) with four poring pulses (20 V, 2 ms pulse length, 50 ms pulse interval, 10% decay rate, positive polarity), followed by five transfer pulses (10 V, 50 ms pulse length, 50 ms pulse interval, 40% decay rate, alternating polarity). After RNP delivery, embryos were washed and cultured in KSOM medium at 37.5 °C under 5% $CO_2$.

**Preparation of donor somatic cells**

Sertoli cells were collected from testes of 1- to 5-day-old 129/Sv × ICR F1 male mice essentially as described previously (*23, 43*). Testes were dissected, decapsulated, and incubated in PBS containing 0.1 mg/mL type IV collagenase (Thermo Fisher Scientific) for 30 min at 37 °C, followed by treatment with 0.25% trypsin and 1 mM EDTA (Thermo Fisher Scientific) for 5 min at room temperature. After four washes with PBS containing 3 mg/mL bovine serum albumin, dissociated cells were resuspended in HEPES-buffered KSOM and used immediately as nuclear donor cells for SCNT.

Peripheral blood donor cells were prepared rom adult male 129/Sv × ICR F1 mice essentially as described previously (*27*). Peripheral blood (~45 µL) was collected from the tail vein using a heparinized calibrated pipette and transferred into a tube containing EDTA. Blood samples were treated with erythrocyte-lysing buffer (155 mM $NH_4Cl$, 10 mM $KHCO_3$, 2 mM EDTA, pH 7.2) for 5 min and washed three times with PBS by centrifugation (1200 × g for 5 min). The resulting leukocyte-enriched cell pellet was resuspended in HEPES-buffered KSOM and used immediately as nuclear donor cells for SCNT. Alternatively, for experiments using cryopreserved donor cells, the isolated leukocytes were resuspended in CELLBANKER 2 (Nippon Zenyaku Kogyo, Fukushima, Japan), transferred to cryovials, and stored at −80°C according to the manufacturer's instructions. After storage for 1 or 30 days, samples were rapidly thawed at 37°C, washed three times with HEPES-buffered KSOM to remove the cryopreservation medium, and immediately used as nuclear donor cells for SCNT.

Tail-tip fibroblasts (TTFs) were established by explant culture from tail biopsies of adult male 129/Sv × ICR F1 mice. Tail tips were minced and cultured in DMEM supplemented with 10% fetal bovine serum and penicillin/streptomycin at 37 °C in 5% $CO_2$. Prior to use as donor cells for SCNT, TTFs were cultured under confluent conditions for an additional 3–4 days to synchronize the cell cycle predominantly at the G0/G1 phase.

**Somatic cell nuclear transfer**

Somatic cell nuclear transfer (SCNT) was performed essentially as described previously with minor modifications (*24, 43, 44*). All recipient oocytes used for SCNT were obtained from superovulated BDF1 females. Enucleation was carried out in the presence of 7.5 µg/mL cytochalasin B (Calbiochem) using a piezo-driven micromanipulator. For SCNT using Sertoli cells or leukocytes, the donor nucleus was injected into the cytoplasm of each enucleated oocyte using a piezo-assisted micromanipulator. For SCNT using TTFs, membrane-intact donor cells were fused with the enucleated oocytes by inactivated Sendai virus envelope (HVJ-E; Ishihara Sangyo). Reconstructed oocytes were activated by incubation in calcium-free KSOM containing 3 mM strontium chloride, 5 µM latrunculin A (LatA; Merck), 50 nM trichostatin A (TSA; Merck), and 1 µM RK701 (MedChemExpress, #HY-152775), a G9a histone methyltransferase inhibitor (G9ai). After 1 hour activation, the embryos were washed and cultured in KSOM (containing calcium) with 5 µM LatA, 50 nM TSA, and 1 µM G9ai. At 8 h post activation (hpa), LatA and TSA were removed, and embryos were further cultured in KSOM containing G9ai alone until 24 hpa. Embryos were then transferred to KSOM. Y-CUT in SCNT embryos was performed by electroporation at 8 hpa using Cas9 ribonucleoprotein complexes targeting the Y chromosome centromere, under the same conditions as those used for Y-CUT in zygotes.

**Embryo transfer**

Embryos that developed to the 2-cell stage were selected and transferred into the oviducts of pseudopregnant ICR females at 0.5 days post coitum (dpc). Pups were delivered by caesarean section at E19.5, examined for external genitalia, and fostered by lactating ICR females.

**DNA-FISH**

The zona pellucida of embryos derived from ICR × ICR crosses were removed by acidic Tyrode's solution at 37°C, and embryos were transferred to M2 medium. Zona-free embryos or cultured cells were fixed in 4% PFA and 0.2% Triton X-100 in PBS for 15 min at room temperature. After washed briefly three times with PBSt (0.05% Triton X-100 in PBS), embryos were permeabilized with 2nd permeabilization buffer (0.5% Triton X-100 and 0.5 mg/mL RNase A in PBS) at 37°C for 1 h. Embryos were then washed two times with PBSt, incubated in 0.1 M HCl containing 0.7% Triton X-100 for 1 min, washed once with PBSt, and incubated in 2× SSC containing 0.1% BSA for 15 min at room temperature. Embryos were then transferred into pre-hybridization buffer (20% formamide, 2× SSC, 0.5 mM EDTA, 10% dextran sulfate, and 0.2% BSA) and incubated for 10 min at room temperature. Embryos were then transferred into hybridization buffer containing FISH probes and equilibrated for 1 h at 37°C. After the incubation, samples were denatured at 80°C for 15 min and subsequently incubated overnight at 37°C in a humidified chamber. Embryos were washed twice with a buffer composed of 2× SSC and 0.01% Triton X-100 for 15 min at 37 °C and once with a buffer composed of 0.4× SSC and 0.01% Triton X-100 for 15 min at 37°C. Embryos were mounted with Vectashield containing DAPI (Vector Labs). For Y chromosome visualization, Y chromosome painting probe (MetaSystems, D-1421-050-FI) was used. X chromosome probe targeting a X chromosome-specific repetitive site was prepared by PCR. A plasmid in which 5'-CAGCTGTGGGTAAGGAAGC-3'sequences were tandemly repeated for 13 times was synthesized and used as a template. PCR products were labelled by Aminoallyl-dUTP-XX-AZDye555 (Jena Bioscience).

**Karyotyping by multicolor FISH**

The spleen was removed from each mouse and minced with scissors, followed by filtration through a 50-µm filter. Each cell suspension was cultured in RPMI 1640 medium supplemented with 10 µg/mL lipopolysaccharide, 3 µg/mL concanavalin A, 50 µM 2-mercaptoethanol, 100 U/mL penicillin, 100 µg/mL streptomycin, and 6 % fetal bovine serum at a density of 2–5×$10^6$ cells/mL for 24–48 h. After the culture, colcemid was added at a final concentration of 0.02 µg/mL and the cells were incubated for 1.5 h. The cells were pelleted by centrifugation at 420 × g for 5 min and resuspended in a hypotonic solution (0.075 M KCl) for 20 min. After hypotonic treatment, cells were initially fixed by adding 1 mL of fixative (methanol:acetic acid = 3:1 v/v), and the tube was immediately and vigorously inverted several times. Subsequently, 5 ml of fixative was added, and the tube was vigorously inverted again. The cell suspension was centrifuged at 420 × g for 5 min, and the supernatant was replaced with 5ml of fresh fixative. This washing step was repeated three times. After washing, chromosome slides were prepared using a Hanabi metaphase spreader (ADSTEC) to both drop 20 µL of the cell suspension onto glass slides and dry them.

For multicolor FISH, the chromosome slides were incubated at 50 °C in air for 2–3 h, and then immersed in 2× SSC in a Coplin jar at 72 °C for 30 min. The Coplin jar was allowed to cool to room temperature for 20 min. The slides were treated in 0.1× SSC for 1 min, followed by denaturation in 0.02 N NaOH at room temperature for 1 min. The slides were then washed in 0.1× SSC and 2× SSC for 1 min each at 4 °C, dehydrated through an ethanol series (70%, 95%, 100%), and air-dried. The hybridization probe (21XMouse; MetaSystems Probes) was denatured at 75 °C for 5 min, cooled in ice water for 30 sec, and pre-incubated at 37 °C for 30 min. The probe was applied onto the slides and incubated for 2 days at 37 °C in a humidified chamber. After hybridization, the slides were treated in 0.4× SSC for 2 min, washed with 2× SSC containing 0.05% Tween-20 for 30 sec, and rinsed with water for 1 min. After drying the slides, they were mounted with DAPI/Antifade (MetaSystems Probes). Fluorescent images were captured using a fluorescence microscope (BX51, Olympus) equipped with a DP75 camera (EVIDENT) and analyzed using ChromaWizard software (*45*). Ten metaphase spreads per mouse were karyotyped to determine the chromosome constitution.

**Quantitative real-time polymerase chain reaction (qPCR)**

Chromosome copy number estimation was carried out as described previously with small modifications (*36, 46*). Genomic DNA was extracted from mouse tail tips and purified by phenol/chloroform extraction followed by ethanol precipitation. The purified DNA was diluted to a concentration of 10 ng/µL and used as the sample. Primer pairs targeting the *Obp1a* and *Sry* genes were used to estimate the copy numbers of the X and Y chromosomes, respectively. Primer pairs for the *Omp* gene located on chromosome 7 and the *Olfr16* gene on chromosome 1 were used for normalization. Two technical replicates were prepared for each reaction, and average values were calculated. The values normalized to each reference gene were then calculated to obtain the final copy number estimate. Genomic DNA from male mice of the same strain was used as a control. qPCR was performed using Thunderbird Next SYBR qPCR Mix (TOYOBO) using a CFX Connect Real-time PCR Detection System (Bio-Rad Laboratories). The primers used for qPCR are as follows.

Olfr16(chr1) -F：GAGTTCGTCTTCCTGGGATTC

Olfr16(chr1) -R：TAATGATGTTGCCAGCCAGA

Omp(chr7)-F：GCCCACTTGATTCCCTGA

Omp(chr7)-R：GCATCTGCTGGGTCAGGTCC

Obp1a(chrX)-F：GGATCAGAATTATGGATCATGTG

Obp1a(chrX)-R：GATCATGAGAAGGGGAAGGA

Sry(chrY)-F：CTCATCGGAGGGCTAAAGTG

Sry(chrY)-R：AAGCTTTGCTGGTTTTTGGA

**Whole-genome sequencing**

Genomic DNA was extracted from the mouse tail tip and purified using the phenol/chloroform extraction method. A total of 1 µg genomic DNA was digested into 200-1000 bp fragments using NEBNext dsDNA Fragmentase (NEB). After digestion, the reaction was terminated by adding 5 µL of 0.5 M EDTA, and DNA fragments were size-selected by removing relatively long DNA with 0.6× AMPure XP (Beckman Coulter). 100 ng of fragmented DNA was resuspended in 20 µl of pure water. The DNA was used for library preparation using NEBnext Ultra II DNA library preparation kit (NEB) following the manufacturer's instruction except that all steps were scaled down to be performed with 40% volume and the samples were subjected to PCR amplification for 8 cycles by KAPA HiFi Hot Start DNA polymerase (Kapa Biosystems) using unique dual-index primers (NEB). DNA was purified by adding 0.8× volume of Ampure XP and eluted in 20 µL of 10 mM Tris-HCl (pH 8.0). The library was validated using TapeStation (Agilent technologies). Paired-end sequencing was performed on an illumina NovaSeq X plus (150 bp × 2).

**Whole-genome sequencing data processing**

Whole-genome sequencing paired-end reads were aligned to the mouse genome (mm39) using bwa-mem2 (version 2.2.1). Properly paired reads were selected, and reads with mismatches were removed using the bamutils filter option with the following parameters: -mismatch 3 -properpair (*47*). Reads from PCR duplicates were removed by using GATK (version 4.6.2.0) (*48*) “MarkDuplicatesSpark” with an option “--remove-all-duplicates true. The DNA copy number was calculated using Control-FREEC (version 11.6) (*49*) with the following parameters: ploidy = 2, breakPointThreshold = .8, window = 1000000, step=500000, sex=XY, mateOrientation = 0, forceGCcontentNormalization = 1. A Merged set of three XY 129×ICR donor samples or an XY ICR samples was used as the control. Read coverage was calculated using mosdepth (version 0.3.11) (*50*). sgRNA target sites including possible off-target sites were identified by Cas-OFFinder (*41*), allowing up to four mismatches. These off-target sites were extended both 100 bp upstream and downstream using bedtools (version 2.31.1) slope (*51*). GATK Haplotypecaller was used to call variants and indel was selected using GATK SelectVariants. GATK Variant Filtration was used to remove variants with the following characteristics QD < 2.0, FS < 200.0, MQ <50.0, DP<10. Variants overlapping XY 129×ICR donor samples were removed using bedtools (version 2.31.1) subtract. Variants overlapping with the putative off-target sites were selected using bcftools (version 1.22) (*52*) and visually validated their accuracy using IGV (version 2.19.7) (*53*).

**Immunostaining**

Fibroblasts were isolated from the mouse tail tips and cultured in DMEM (Nacalai tesque) supplemented with 10% FBS and penicillin/streptomycin. Fibroblasts were fixed with 4% paraformaldehyde in PBS for 10 min at room temperature, followed by permeabilization with 0.2% Triton X-100 in PBS for 10 min at room temperature. Blocking was performed with a blocking buffer (3% BSA in PBS) for 30 min. Fibroblasts were then incubated with anti-H3K27me3 antibody (Cell Signaling Technology #9733, 1:1000) for 1 h at room temperature. After washing three times with 0.02% Triton X-100 in PBS, the cells were incubated with a secondary antibody, donkey Alexa Fluor 555-conjugated anti-rabbit IgG (Thermo Fisher #A-31572; 1:1000), for 1 h at room temperature. Samples were mounted with VECTASHIELD and imaged using an FV1200 confocal microscope (Olympus).

**Statistical analysis**

Statistical analyses were implemented with GraphPad Prism or R software (http://www.r-project.org).

### Fig. S1

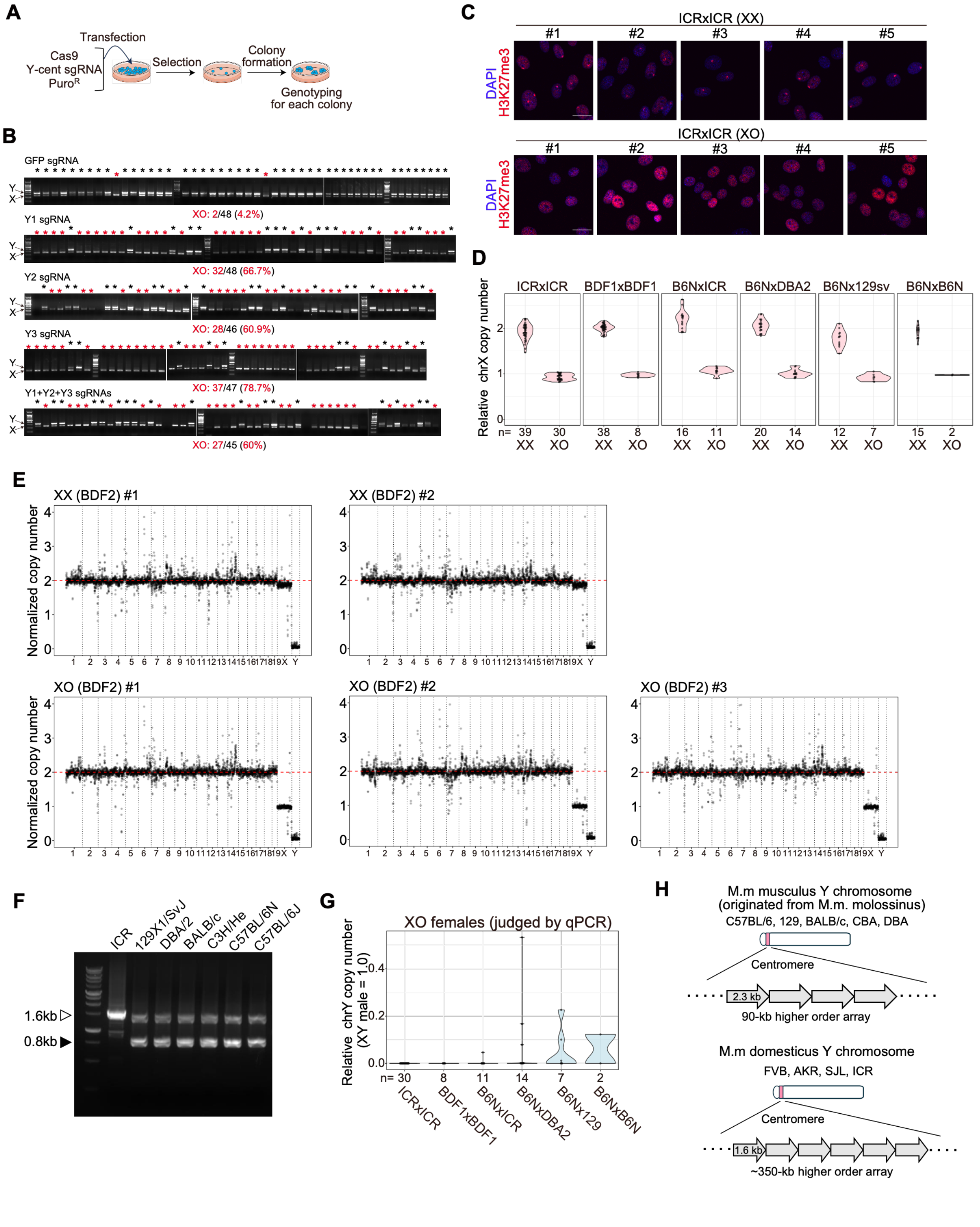
A
Transfection
Cas9
Y-cent sgRNA
PuroR
Selection
Colony formation
Genotyping for each colony
B
GFP sgRNA
XO: 2/48 (4.2%)
Y1 sgRNA
XO: 32/48 (66.7%)
Y2 sgRNA
XO: 28/46 (60.9%)
Y3 sgRNA
XO: 37/47 (78.7%)
Y1+Y2+Y3 sgRNAs
XO: 27/45 (60%)
C
ICRxICR (XX)
ICRxICR (XO)
#1
#2
#3
#4
#5
DAPI
H3K27me3
D
Relative chrX copy number
ICRxICR
BDF1xBDF1
B6NxICR
B6NxDBA2
B6Nx129sv
B6NxB6N
n= 39 30 38 8 16 11 20 14 12 7 15 2
XX XO
E
XX (BDF2) #1
XX (BDF2) #2
XO (BDF2) #1
XO (BDF2) #2
XO (BDF2) #3
Normalized copy number
F
ICR
129X1/SvJ
DBA/2
BALB/c
C3H/He
C57BL/6N
C57BL/6J
1.6kb
0.8kb
G
XO females (judged by qPCR)
Relative chrY copy number (XY male = 1.0)
n= 30 8 11 14 7 2
ICRxICR
BDF1xBDF1
B6NxICR
B6NxDBA2
B6Nx129
B6NxB6N
H
M.m musculus Y chromosome
(originated from M.m. molossinus)
C57BL/6, 129, BALB/c, CBA, DBA
Centromere
2.3 kb
90-kb higher order array
M.m domesticus Y chromosome
FVB, AKR, SJL, ICR
Centromere
1.6 kb
~350-kb higher order array

**Fig. S1. Targeted elimination of the Y chromosome in cultured cells and mice**

**(A)** Experimental scheme to assess Y chromosome elimination by Y-CUT in mESCs.
**(B)** Agarose gel electrophoresis images. PCR was performed to examine the presence of X and Y chromosomes. Y1, Y2, or Y3 sgRNA were expressed in mESCs together with Cas9. sgRNA against GFP was used as a negative control.
**(C)** Immunofluorescence images of H3K27me3. H3K27me3 staining on mouse tail fibroblasts was performed to confirm the XX or XO status. H3K27me3 spots in XX cells indicate the presence of inactivated X chromosome. Scale bars, 20 µm.
**(D)** Violin plots showing copy numbers of the X chromosome in indicated samples. Chromosome copy numbers were determined by qPCR.
**(E)** Plots showing the chromosome copy numbers. Genome sequencing coverages at 1-Mb bins were plotted. Data was normalized to the XY genome sequencing data.
**(F)** Agarose gel electrophoresis image. PCR was performed to examine the origin of the Y chromosomes in indicated mouse strains. Amplification of 0.8 and 1.6 kb bands indicates the presence of the *M.m musculus*-type Y chromosome and the *M.m domesticus*-type Y chromosome, respectively.
**(G)** Violin plots showing relative copy numbers of the Y chromosome in indicated samples. All the samples contained single X chromosome. Chromosome copy numbers were determined by qPCR.
**(H)** Schematic illustration indicating the structural difference of the Y chromosome centromeres among mouse strains. The *M.m domesticus*-type Y chromosome carries larger centromere with shorter array units compared to the *M.m musculus*-type Y chromosome.

#### Fig. S2

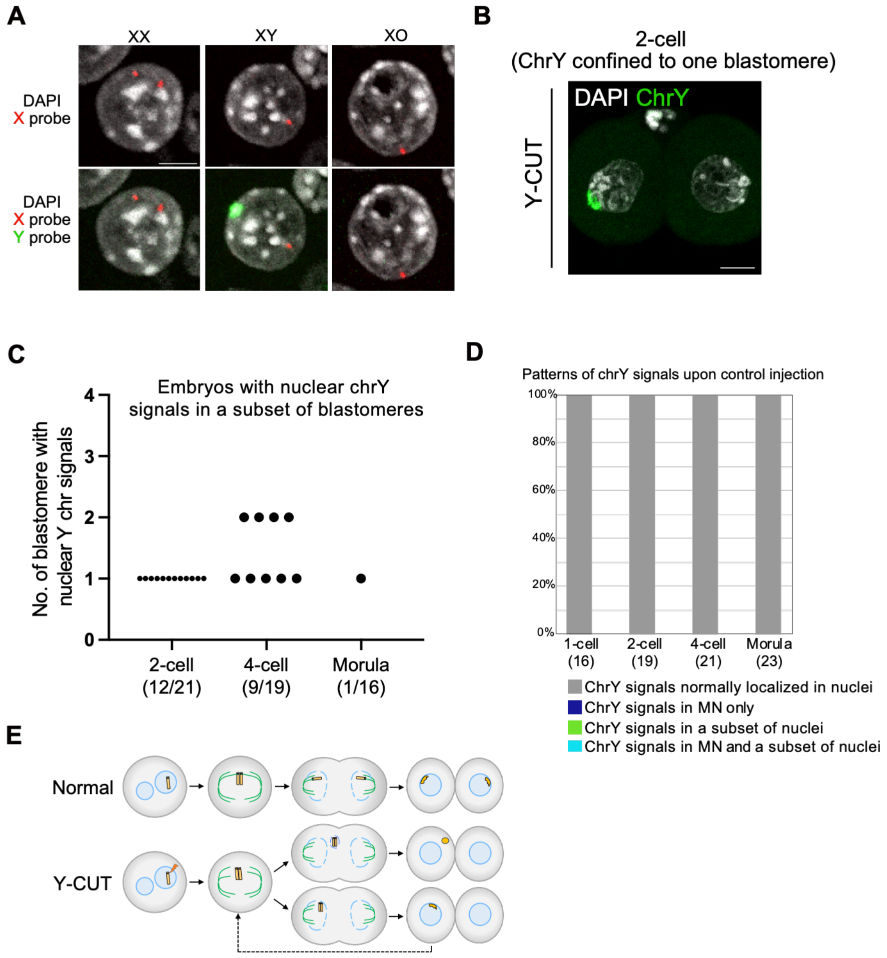


**Fig. S2. Y chromosome is released as micronuclei upon Y-CUT during early development**

**(A)** Representative DNA-FISH images in mouse embryos. DNA-FISH probes for the X and Y chromosomes were used to discriminate XX, XY, and XO statuses. Scale bars, 5 µm.

**(B)** Representative image of the Y chromosome retained only in one blastomere at the 2-cell stage after Y-CUT. Scale bar, 10 µm.

**(C)** Plots showing the numbers of blastomeres showing nuclear Y chromosome signals at the indicated stages after Y-CUT.

**(D)** Stacked bar graph showing the frequency of the embryos showing the indicated Y chromosome signal localization patterns. The graph indicates the results in control embryos.

**(E)** Schematic illustration showing the expected outcome of Y-CUT in zygotes. Centromere-inactivated Y chromosomes tend to localize outside of nuclei after cell divisions due to chromosome missegregation and eventually form micronuclei.

#### Fig. S3

**A**

| Donor | | | crY3* | No. of total ET | No. of implant (% per ET) | No. of pup (% per ET) | No. of male (% per total) | No. of female (% per total) |
|---|---|---|---|---|---|---|---|---|
| Cell type | Age | Freeze (day) | | | | | | |
| Sertoli | 2-3 days | - | - | 45 | 23 (51.1) | 1 (2.2) | 1 (100) | 0 (0) |
| | | - | + | 118 | 57 (48.3) | 13 (11.0) | 3 (23) | 10 (77) |
| TTF | 8-11 weeks | - | - | 14 | 9 (64.3) | 3 (21.4) | 3 (100) | 0 (0) |
| | | - | + | 26 | 18 (69.2) | 4 (15.4) | 0 (0) | 4 (100) |
| Blood | 8-11 weeks | - | - | 88 | 47 (53.4) | 7 (8.0) | 7 (100) | 0 (0) |
| | | - | + | 143 | 71 (49.7) | 11 (7.7) | 1 (9) | 10 (91) |
| | | 1 | - | 43 | 23 (53.5) | 6 (14.0) | 6 (100) | 0 (0) |
| | | 1 | + | 35 | 17 (48.6) | 2 (5.7) | 0 (0) | 2 (100) |
| | | 30 | - | 60 | 27 (45.0) | 3 (5.0) | 3 (100) | 0 (0) |
| | | 30 | + | 128 | 63 (49.2) | 5 (3.9) | 0 (0) | 5 (100) |

*crY3 was introduced by electroporation together with tracrRNA and spCas9 protein.
TTF, tail tip fibroblast. ET, embryo transfer.

**B**

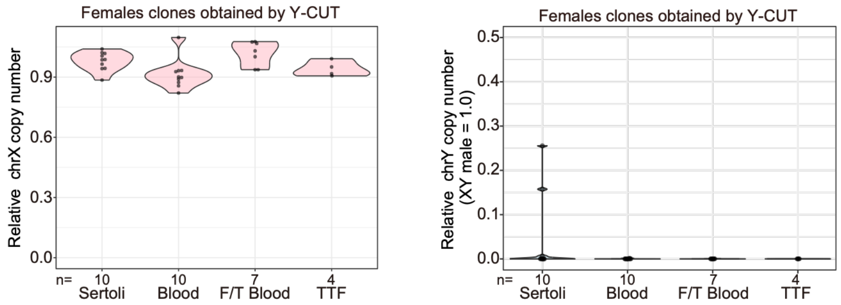


**C**

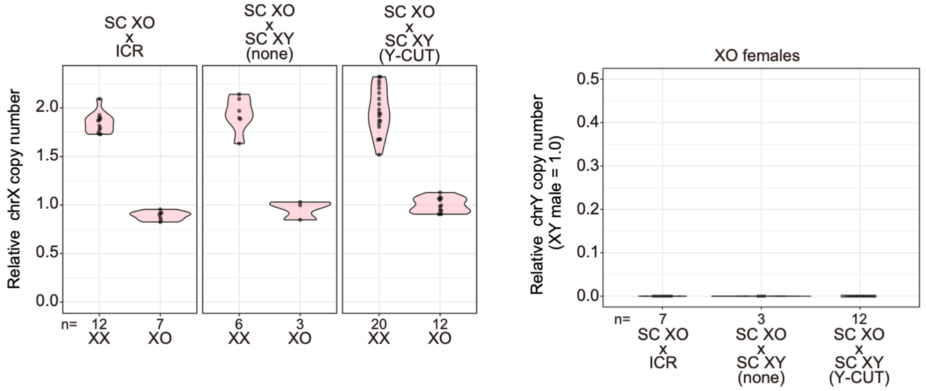


**Fig. S3 Characterization of female clones generated by dual-sex cloning**
**(A)** Table showing the birth rates and sexes of cloned pups after somatic cell nuclear transfer.
**(B)** Violin plots showing copy numbers of the X chromosome (left) and Y chromosome (right) in indicated XO female samples. Chromosome copy numbers were determined by qPCR. F/T, freeze/thaw.
**(C)** Violin plots showing copy numbers of the X chromosome (left) and Y chromosome (right) in offspring from indicated crosses. Those crosses generated XX and XO females. Chromosome copy numbers were determined by qPCR. SC XO, Sertoli cell nuclear transfer-derived XO females generated through Y-CUT; SC XY, Sertoli cell nuclear transfer-derived XY males.

#### Fig. S4

**A**

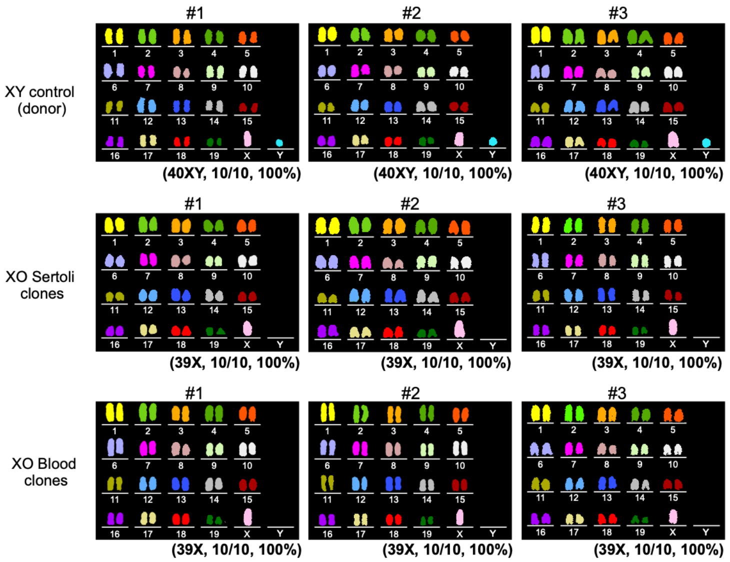


**B**

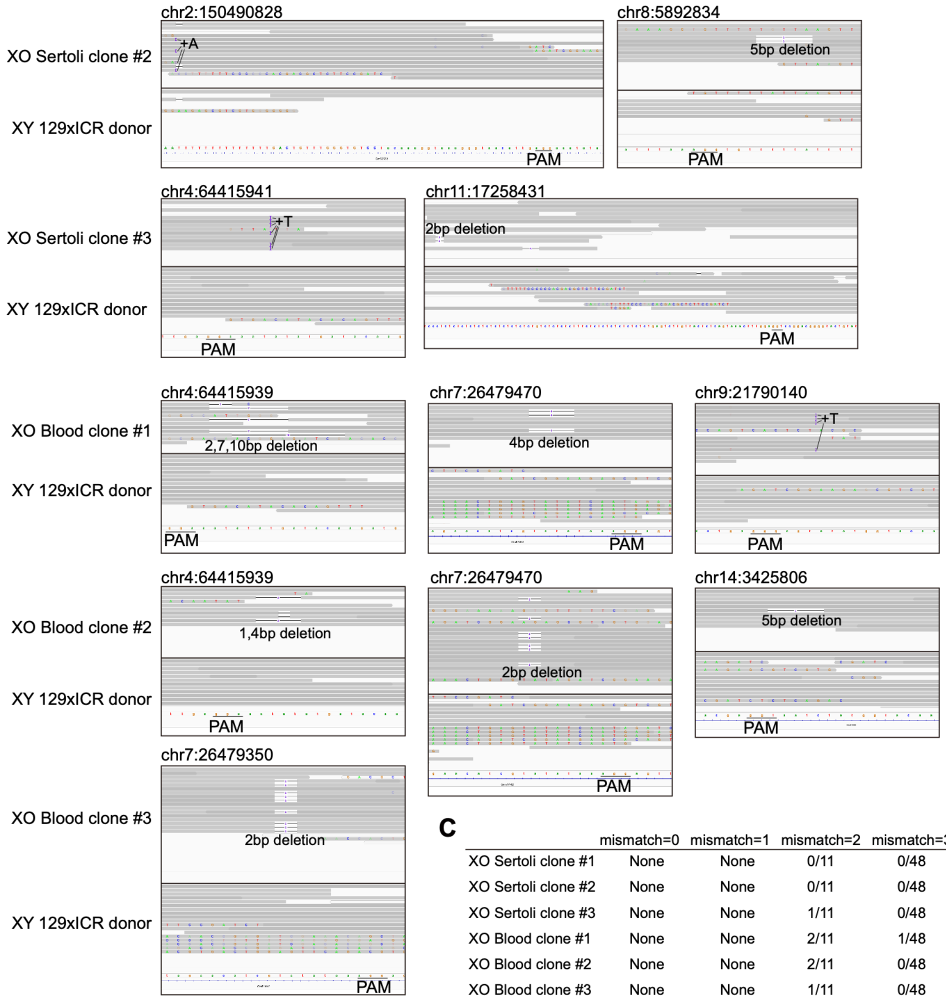


**C**

| | mismatch=0 | mismatch=1 | mismatch=2 | mismatch=3 | mismatch=4 |
|---|---|---|---|---|---|
| XO Sertoli clone #1 | None | None | 0/11 | 0/48 | 0/270 |
| XO Sertoli clone #2 | None | None | 0/11 | 0/48 | 2/270 |
| XO Sertoli clone #3 | None | None | 1/11 | 0/48 | 1/270 |
| XO Blood clone #1 | None | None | 2/11 | 1/48 | 0/270 |
| XO Blood clone #2 | None | None | 2/11 | 0/48 | 1/270 |
| XO Blood clone #3 | None | None | 1/11 | 0/48 | 0/270 |

**Fig. S4. Genomic analyses of female clones generated by dual-sex cloning**

**(A)** Karyotypes of male donor mice and cloned female mice generated through Y-CUT. Multicolor FISH was performed to examine their karyotype. Three independent individuals (#1 to #3) were analyzed. All ten examined somatic cells in donor mice showed normal karyotype (40XY), while those in Sertoli cell- or male blood cell-derived cloned female mice generated through Y-CUT indicated XO (39X) karyotype. No Robertsonian or reciprocal translocations were observed except for the complete loss of the Y chromosome in the cloned female mice generated by Y-CUT.

**(B)** Examination of off-target effects of Y-CUT. Detected nucleotide alterations at the possible off-target sites are indicated. Only a few small indels were observed.

**(C)** Table showing the frequency of the indel occurrence at the possible off-target sites.

#### Fig. S5

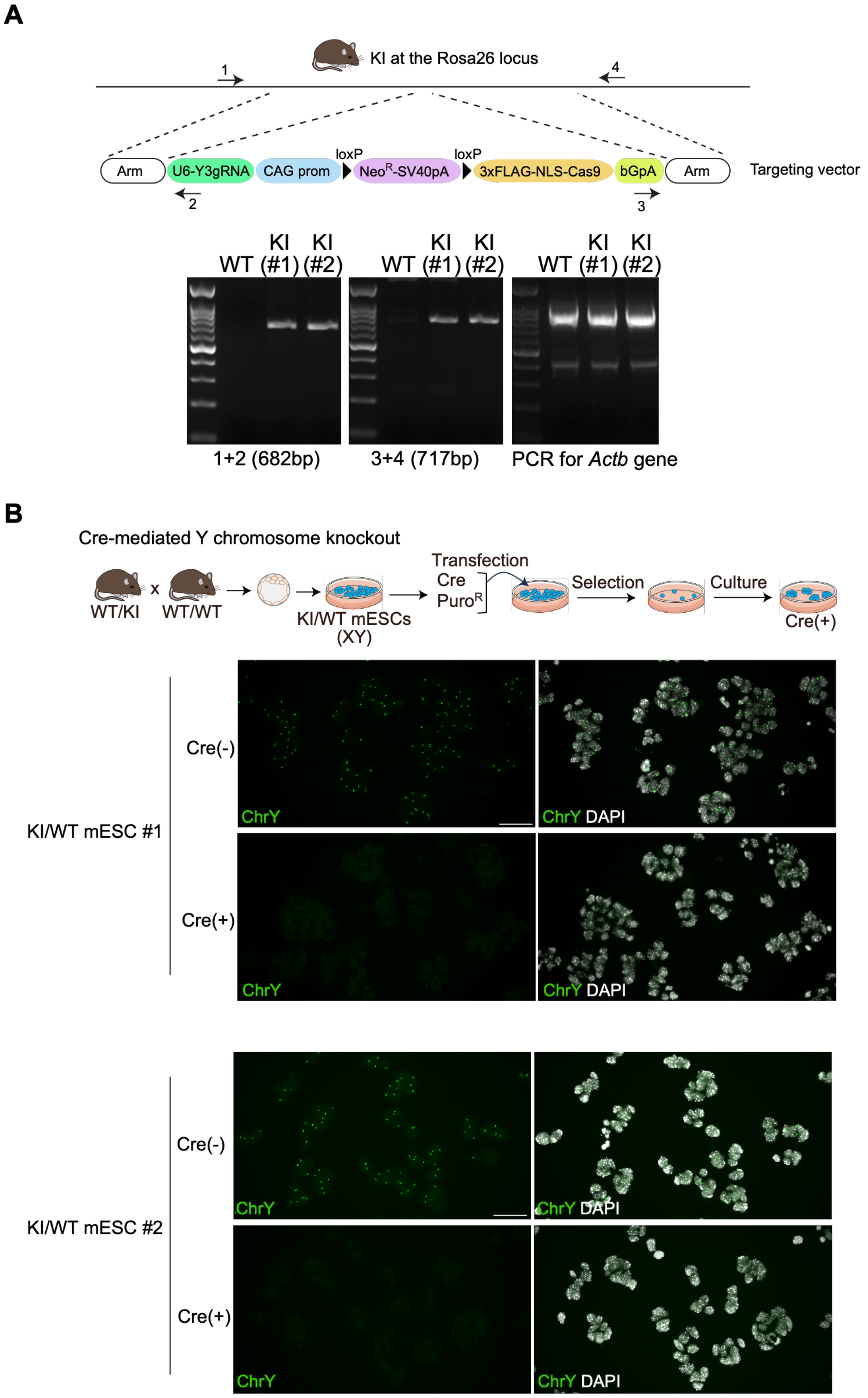
A
KI at the Rosa26 locus
1
4
Arm
U6-Y3gRNA
CAG prom
loxP
NeoR-SV40pA
loxP
3xFLAG-NLS-Cas9
bGpA
Arm
Targeting vector
2
3
WT KI (#1) KI (#2)
WT KI (#1) KI (#2)
WT KI (#1) KI (#2)
1+2 (682bp)
3+4 (717bp)
PCR for Actb gene
B
Cre-mediated Y chromosome knockout
WT/KI x WT/WT
KI/WT mESCs (XY)
Transfection
Cre
PuroR
Selection
Culture
Cre(+)
KI/WT mESC #1
Cre(-)
Cre(+)
ChrY
ChrY DAPI
KI/WT mESC #2
Cre(-)
Cre(+)
ChrY
ChrY DAPI

**Fig. S5. Genetic encoding of Y-CUT**

**(A)** Generation of knock-in mice carrying a genetically encoded Y-CUT cassette. The agarose gel images show genotyping PCR results using indicated primers (arrows with numbers).

**(B)** Top, experimental scheme for Cre-mediated Y chromosome knockout in mESCs. Heterozygous knock-in (KI/WT) mESCs carrying the genetically encoded Y-CUT cassette was derived from blastocysts generated by the cross between heterozygous KI mice and wild-type mice. Cre was then transiently expressed in KI/+ mESCs, and Cre-expressing cells were selected by puromycin. Bottom, DNA-FISH for the Y chromosome was performed with or without Cre expression. Scale bars, 50 µm.

Type or paste caption here. Create a page break and paste in the figure above the caption.

**Table S1. List of sgRNAs targeting the Y chromosome centromere**

**Table S2. Specificity of sgRNAs used in this study**

**Movie S1. Three-dimensional reconstruction of confocal DNA-FISH images of 2-cell embryos.**

**Movie S2. Three-dimensional reconstruction of confocal DNA-FISH images of 4-cell embryos.**

**Movie S3. Three-dimensional reconstruction of confocal DNA-FISH images of morulae**
